# GPR97/ADGRG3 is activated by its tethered peptide agonist and not by steroids to induce neutrophil polarization and migration

**DOI:** 10.1101/2025.09.11.675646

**Authors:** Tyler Bernadyn, Frank Kwarcinski, Naincy Chandan, Riya Gandhi, Yuling Feng, Michael Holinstat, Carole A. Parent, Alan V. Smrcka, Gregory G. Tall

## Abstract

Most adhesion G protein coupled receptors (AGPCRs) are activated by intramolecular binding of a tethered-peptide agonist (TA). Shear force-induced dissociation of the AGPCR N-terminal fragment (NTF) and C-terminal fragment (CTF) exposes the TA. The decrypted TA binds rapidly to its orthosteric site within the CTF to stabilize the active state of the AGPCR. Corticosteroids were previously proposed to be agonists for GPR97/ADGRG3. Later, other steroids and androgens were purported to be selective agonists of additional AGPCRs. Here, we demonstrate that GPR97/ADGRG3 is activated by dissociation of its NTF/CTF and follows the TA mechanism. TA peptidomimetics and the ADGRG subfamily partial agonist 3-acetoxydihydrodeoxygeduin (3-α-DOG), but not corticoids, stimulated GPR97/ADGRG3 in cell-based luciferase reporter assays and receptor/G protein reconstitution assays. GPR97 was defined as a promiscuous AGPCR that couples to G13, Gs and Gi, but not Gq. GPR97 is highly expressed in human polymorphonuclear neutrophils (hPMNs). Neutrophils undergo actin polymerization-induced cell shape changes and polarization and migrate upon activation. We found that GPR97 activation via TA peptidomimetics or 3-α-DOG robustly stimulated hPMN and mouse bone-marrow neutrophil (mBMN) polarization. Furthermore, GPR97 TA peptidomimetics and 3-α-DOG, but not beclomethasone, induced hPMN and mBMN chemotaxis. Together, our results demonstrate that GPR97/ADGRG3 utilizes a tethered agonist mechanism to activate G protein signaling and induce neutrophil polarization and migration.

**One Sentence Summary:** GPR97/ADGRG3 tethered agonism regulates G protein signaling in neutrophils.

## INTRODUCTION

G protein coupled receptors (GPCRs) are the most prevalent class of membrane receptors and pharmaceutical targets (1). Class B2 or adhesion GPCRs (AGPCRs) are the second largest GPCR subfamily consisting of 33 receptors. AGPCRs have important roles in a variety of physiological systems ranging from embryonic development, maintenance of proper neurological function, and regulation of circulating cell dynamics (2–4). AGPCRs consist of N-terminal ectodomains made up of a variety of adhesive sub-domains and a common GPCR autoproteolysis-inducing (GAIN) domain (5, 6). Most AGPCRs are orphans that lack identified ligands that bind to the ectodomains. The GAIN domain is positioned at the C-terminal end of the ectodomain and has two subdomains including a variable length α-helical-abundant GAIN_A_ subdomain and a conserved GAIN_B_ subdomain with a β-sheet core. Most AGPCRs undergo self-proteolysis at the GPCR proteolytic site (GPS) which is situated between the penultimate and last β-strands of GAIN_B_. The self-cleavage reaction is thought to be constitutive and to occur during receptor biosynthesis such that the resultant two-fragment receptor, composed of an N-terminal fragment (NTF) and a C-terminal fragment (CTF) or the 7TM domain arrives at the plasma membrane in a noncovalently-bound topology (6). In this holoreceptor form, the last β-strand remains embedded within the core of the GAIN domain. Ligand binding anchors the NTF and cell movement opposed to this anchor is thought to dissociate the NTF from the CTF, which pulls the β-strand out of the GAIN domain where it undergoes a conformation change into a partial α-helix as or after it enters the CTF/7TM orthosteric site to become a tethered-peptide agonist (TA) (7, 8). A majority of AGPCRs are thought to utilize this mechanism of activation; however, others may follow the proposed “tunable” model in which the intact holoreceptor may undergo subtle conformational changes in response to ligand binding that results in G protein signaling (9, 10).

All members of the AGPCR ADGR’G’ subfamily that have been tested follow the TA activation mechanism (7, 8, 10, 11). However, GPR97/ADGRG3 was proposed to be activated by beclomethasone and cortisol, steroids that may serve as diffusible agonists within the orthosteric site of the receptor in lieu of the TA (12). More recently, additional steroids were proposed to activate additional AGPCRs with surprising exclusivity and receptor specificity. Dehydroepiandrosterone (DHEA) activated GPR64 (13). 5-α-dihydrotesterone (5α-DHT) activated GPR133 to increase cAMP levels in muscle explants (14). 17α-hydroxypregnenolone was purported to activate GPR56 to prevent liver injury (15). The four ring backbone of steroids is rich in hydrocarbons and amenable to residing in hydrophobic spaces such as the core of GPCR 7TM barrels. The AGPCR orthosteric site is a large hydrophobic cavity that is comprised of conserved residues that evolved to accommodate the seven amino acid TA, which consists of the highly hydrophobic consensus sequence, TϕFϕϕLM where ‘ϕ’ represents any hydrophobic residue (5, 7, 11, 16). The most critical TA residues are also the most conserved, the P3 phenylalanine, the P6 leucine, and the P7 methionine are arranged via a hook-like conformation to reside deepest within the hydrophobic orthosteric site. This hook-like, partial α-helical turn configuration of the TA is common in all TA-activated AGPCR solved structures (7, 8, 11, 17, 18).

Three of the ADGR’G’ subfamily receptors GPR56 (G1), GPR97 (G3), and GPR114 (G5) appear to be the products of gene duplication and are present on three different hematopoietic stem cell lineages, platelets, neutrophils, and eosinophils, respectively (2, 19). Neutrophils are part of the innate immune system and provide the first line of defense at sites of infection and injury (20, 21). To reach these sites, they cross blood vessels via trans-endothelial migration (TEM), a process that involves Gi-mediated adhesion, polarization and migration (22). In addition, neutrophil polarization is also influenced by G13 signaling, which activates Rho small GTPases to induce actin polymerization and cell shape changes (20, 23). Following polarization, neutrophils extravasate through the endothelial cells and chemotax towards infected/injured sites where they contain the affected area by phagocytosing pathogens and releasing powerful proteases, cytokines and neutrophil extracellular traps (NETs) (20). Chemotactic GPCRs that activate Gi signaling are well described, but the GPCRs that may activate G13 signaling during neutrophil TEM are not well characterized. A close homolog of GPR97, GPR56 was shown to facilitate shear-force and collagen-dependent platelet shape change and filipodia protrusion by activating G13 signaling (2, 10, 24). Therefore, we investigated GPR97 and its potential to activate G13 signaling to regulate neutrophil shape change, polarization, and migration. Given that select steroids were proposed as agonists for GPR97, we first provide a detailed analysis to show that GPR97 is activated by its TA and then performed a direct, quantifiable comparison of receptor efficacy imparted by the TA or steroids. We also report that treatment of isolated human or mouse neutrophils with GPR97 TA peptidomimetics, or the ADGRG partial agonist 3-acetoxydihydrodeoxygeduin (3-α-DOG), but not by steroids, induced neutrophil polarization and migration.

## RESULTS

### GPR97 activates G13, Gs, Gi, but not Gq

The specificity of GPR97 G protein coupling has been enigmatic. One Gαq/o chimeric protein of a few tested was activated by GPR97 in a cell-based assay, and a structure of GPR97 bound to mini-Go was solved (12, 25). However, the receptor complex was stabilized by a palmitate that was attached to mini-Gαo and made strong contacts within the core of the GPR97 7TM barrel (12, 17). Endogenous Gαo has not been shown to be modified with palmitate at Cys 351, but this residue can be modified by *Pertussis* toxin. To comparably assess GPR97 coupling to different G proteins, we reconstituted prepared membranes expressing TA-active GPR97-CTF with purified Gβ_1_Gγ_2_ and Gα13, Gαs_s_, Gαq, or Gαi_1_ and measured the kinetics of receptor-stimulated G protein heterotrimer GTPγS binding. Positive receptor membrane controls for G protein activation were TA-active GPR56-CTF (G13) (7), TA-active GPR114-CTF (Gs) (11, 26), TA-active GPR110-CTF (Gq) (18), and the muscarinic acetylcholine receptor 2 (M2R) with carbachol stimulation (Gi_1_) (27). GPR97 showed robust activation of G13 and Gs, and comparable activation of Gi_1_ to M2R, although Gi/o assays are affected by a high constitutive activity present in the empty membrane preparations (Figure 1A-C). GPR97 did not couple to Gq (Figure 1D).

**Figure 1:**
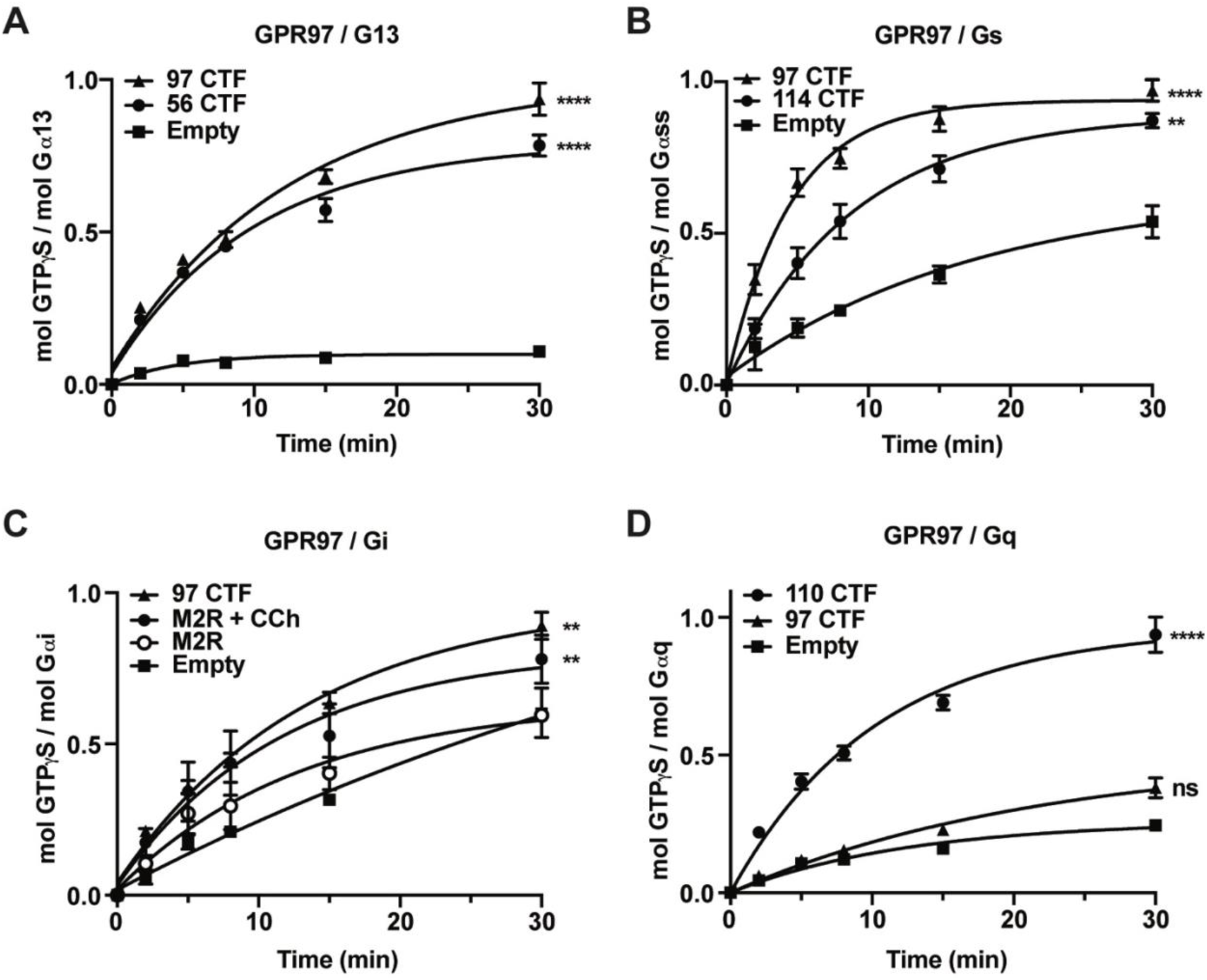
GPR97 couples to G13, Gs, and Gi, but not Gq. GPR97 CTF and empty membranes were tested for stimulation of GTPγS binding kinetics of (**A**) G13 alongside the GPR56 CTF, (**B**) Gs alongside the GPR114 CTF, (**C**) Gi alongside the M2 muscarinic receptor ± carbachol, and (**D**) Gq alongside the GPR110 CTF. Biological triplicate reactions with technical triplicates are presented. Error bars are the means ± S.D. AUCs were calculated using GraphPad Prism and Tukey’s Multiple Comparison tests were used to evaluate differences to the no treatment or empty (no receptor) membrane conditions; ns, not significant, *p < 0.05, ** p < 0.01, *** p < 0.001, **** p < 0.0001.

### GPR97 has a dissociable, glycosylated NTF and is activated by its tethered-peptide-agonist

There has been no demonstration that GPR97 is activated by its tethered agonist (TA) despite its similarity to the other adhesion GPCR ‘G’ subfamily members that are TA-activated (5). We first verified a report that GPR97 is autoproteolytically cleaved in its GAIN domain (19). Membrane samples were prepared from *Sf*9 cells that overexpressed human wild type (WT) GPR97, cleavage-deficient GPR97 (ClvDef) harboring a mutated GPS, and a dual TA mutant (L^P6’^A / L^P7^’A) that for other adhesion GPCRs maintains cleavage but abolishes tethered agonism (28, 29). Membranes with combined ClvDef and dual TA GPR97 mutations were also produced. The GPR97 membrane samples were immunoblotted with 97-NTF (Figure 2A) and 97-CTF (Figure 2B) antibodies. WT and dual TA mutant GPR97 were partially cleaved showing a glycosylated uncleaved band at ∼60 kDa, glycosylated NTF species at ∼32-34 kDa, and a non-glycosylated CTF band at 25 kDa that matched the band observed from membranes expressing GPR97-CTF only (Figure 2B). The predicted molecular weight of the GPR97 CTF is 33.3 kDa; however, adhesion GPCR CTFs migrate aberrantly fast via SDS-PAGE. When the membrane samples were treated with PNGaseF to remove N-linked glycans, deglycosylated, uncleaved GPR97 species were observed at ∼45-48 kDa. A sharp deglycosylated NTF band was observed at ∼26 kDa which aligns with the predicted molecular weight of the GPR97 NTF (Figure 2A). ClvDef and ClvDef plus TA mutant GPR97 only had the glycosylated bands at ∼60 kDa and a deglycosylated species at ∼45-48 kDa. No MW shifts were observed for the GPR97 CTF upon PNGaseF treatment showing that is not modified by N-linked glycans.

**Figure 2:**
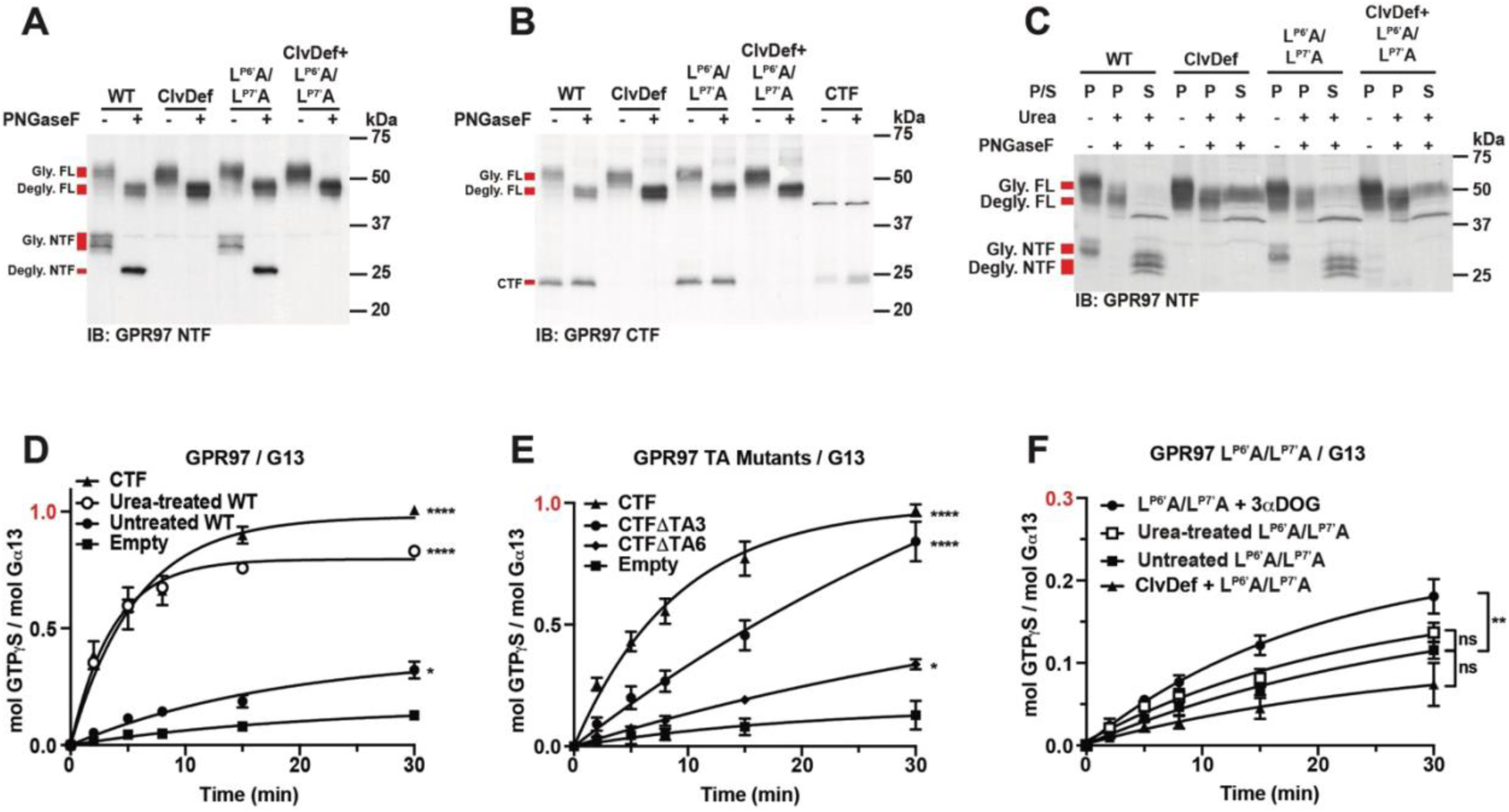
GPR97 is activated by its tethered peptide agonist following NTF/CTF dissociation. Wild type, cleavage deficient mutant (ClvDef), tethered agonist double point mutant (L^P6’^A/L^P7’^A), and cleavage deficient plus tethered agonist double point mutant GPR97 membranes were treated ± PNGaseF and immunoblotted with GPR97 (**A**) NTF-or (**B**) CTF-specific antibodies. (**C**) The GPR97 membranes were treated ± PNGaseF, then ± urea, centrifuged and the urea-soluble (S) and –pellet (P) fractions were immunoblotted with the GPR97 NTF-specific antibody. (**D – F**) Receptor membrane homogenates were mock treated (solid symbols) or urea-treated (open symbols) and reconstituted with purified Gα13 and Gβ_1_Gγ_2_ prior to measurement of G13 GTPγS binding kinetics as follows, (**D**) GPR97 wild type (WT), CTF, or empty membranes, (**E**) GPR97-CTF, –CTΔTA3, –CTFΔTA6, or empty membranes, (**F**) GPR97 tethered agonist double point mutant ± cleavage deficient (ClvDef) mutant membranes were treated ± 25 µM 3-α-DOG as indicated. Biological triplicate reactions with triplicate technical replicates are presented. Error bars are the means ± S.D. AUCs were calculated using GraphPad Prism and Tukey’s Multiple Comparison tests were used to evaluate differences compared to no treatment or empty (no receptor) membrane conditions; ns, not significant, *p < 0.05, ** p < 0.01, *** p < 0.001, **** p < 0.0001.

The GPR97 membranes were then treated with urea to determine if the NTF could be dissociated and solubilized. We used urea before as a proxy for simulating force-induced adhesion GPCR NTF removal and TA-induced receptor activation (10, 29). After urea treatment, membranes were fractionated by centrifugation at 100,000 g to separate membrane pellets (P) from soluble (S) fractions for immunoblotting with the h97NTF antibody. ClvDef receptors largely remained associated with the membrane, whereas the NTFs of WT and TA dual mutant GPR97 were shifted to the urea soluble fraction (Figure 2C).

We investigated the requirements of the TA and NTF dissociation-mediated TA decryption for GPR97 activation of G13. Urea treatment dramatically activated WT GPR97 to enhance G13 GTPγS binding kinetics to match those stimulated by the GPR97 CTF-only receptor (Figure 2D). Non-urea treated WT GPR97 exhibited low activity that was elevated modestly above empty membranes, accounting for receptor basal activity. Two truncations to the TA sequence were made to the GPR97 CTF-only construct that removed the first 3 or 6 TA residues (CTFΔTA3 and CTFΔTA6). A graded diminution of G13 activation was observed as these portions of the TA were truncated (Figure 2E). We further assessed the importance of the TA for receptor activation by showing that dual TA mutant GPR97 (L^P6’^A / L^P7^’A) was not stimulated by urea but was modestly activated by the adhesion GPCR “G” subfamily, small molecule partial agonist, 3-α-DOG (Figure 2F) (30). Urea-mediated NTF dissociation decrypts the defective TA, which does not provide receptor activation, whereas the partial stimulation by 3-α-DOG indicates that dual TA mutant GPR97 retains ability to be activated pharmacologically.

### GPR97 is activated by a tethered agonist peptidomimetic

ADGRG1/GPR56 and ADGRG5/GPR114 are closely related to ADGRG3/GPR97 and can be activated by 7 and 18/19 amino acid synthetic peptides that were designed after the respective receptor TA and stalk sequences (10, 26, 29). To determine whether GPR97 synthetic TA peptidomimetics could activate the receptor in cells, we first calibrated HEK293T cell-based CRE– and SRE-luciferase (LUC) assays to corroborate the finding that GPR97 couples strongly to Gs and G13, respectively (Figure 3A and 3B). Plasmid transfection titrations were performed of the GPR97 CTF with an intact TA and the compromised TA mutants (ΔTA3 and ΔTA6). The GPR97 CTF strongly activated the CRE-LUC reporter, whereas both TA mutants had markedly reduced activities. Interestingly, the GPR97-CTFΔTA3 and ΔTA6 mutants showed the same TA-dependence in the SRE-LUC assay when low amounts of receptor plasmids were transfected, but the pattern became askew with high transfected amounts. We do not fully understand the latter SRE-LUC assay results, because the same mutant receptors exhibit the expected TA-dependence in the Gs and G13 reconstitution assays, as well as the CRE-LUC assay (Figure 3E and 3F, Figure 3A). We speculate that there may be TA-dependent effects on receptor trafficking that account for the differences in the cell-based CRE-LUC and SRE-LUC assays.

**Figure 3:**
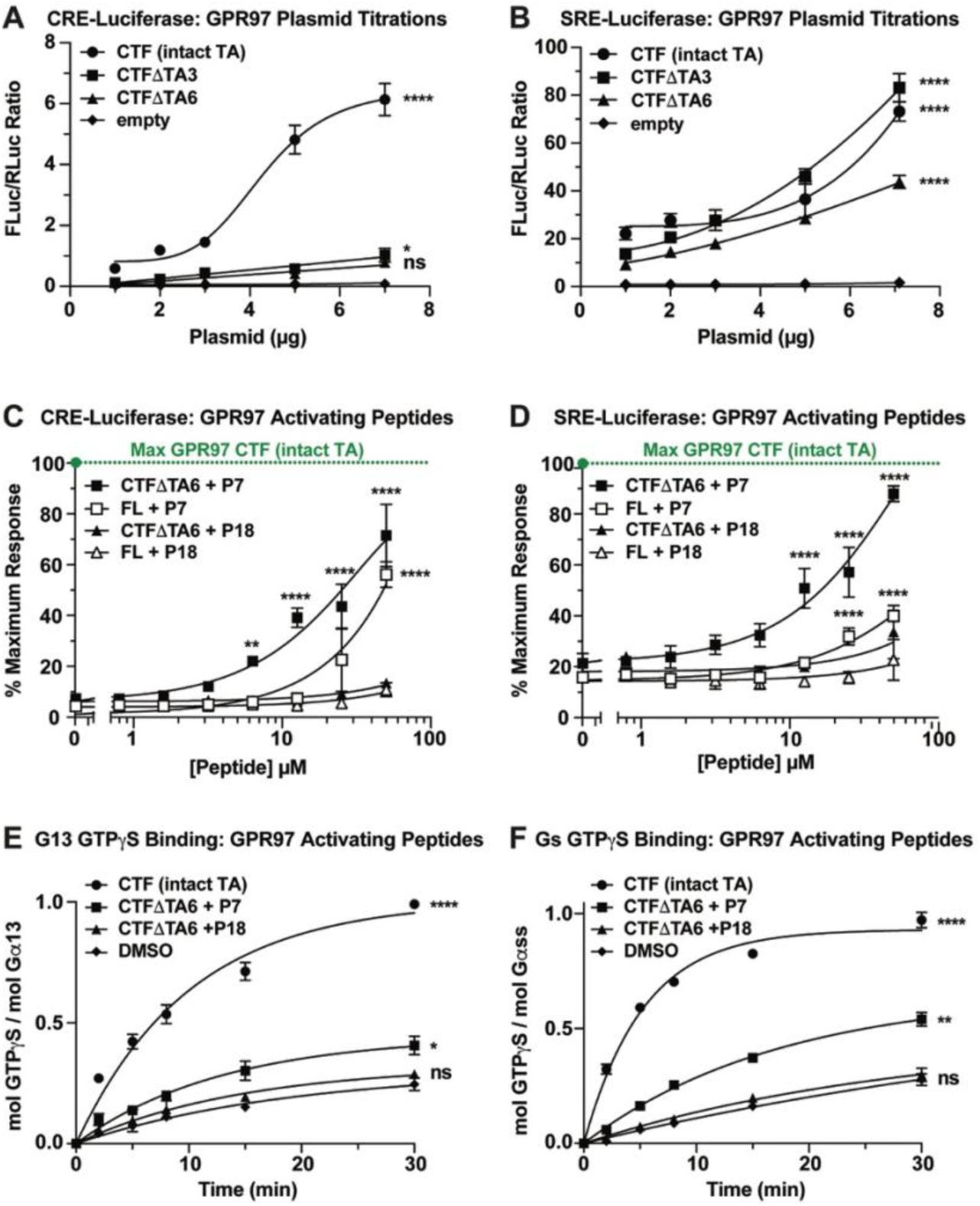
GPR97 tethered agonist-dependent activation of Gs and G13 signaling in cells. (**A**) SRE-luciferase and (**B**) CRE-luciferase activities were measured in HEK293T cells transfected with the indicated amounts of the GPR97-CTF (intact TA) construct, GPR97-CTF constructs with compromised tethered agonists (ΔTA3 or ΔTA6), or empty vector. (**C**) CRE– and (**D**) SRE-luciferase activities of full length (FL) GPR97 and GPR97 CTFΔTA6 via were measured in HEK293T cells in response to the GPR97 tethered agonist peptidomimetics, P7 and P18. The maximal efficacy of GPR97-CTF with an intact TA from (**A**) and (**B**) was denoted by the dashed green lines. (**E**) Kinetics of G13 GTPγS binding stimulated by GPR97-CTF or GPR97-CTFΔTA6 ± tethered agonist peptidomimetics (P7 and P18). (**F**) Kinetics of Gs GTPγS binding stimulated by GPR97 CTF or GPR97-CTFΔTA6 ± tethered agonist peptidomimetics (P7 and P18). Data are the ratios of the FLuc reporter signal to RLuc balancer signal and are biological triplicates. **C** and **D** data was presented as percent of the maximal GPR97-CTF (intact TA) signal and two-way ANOVAs were used for statistical analyses. AUCs were calculated for the **E** and **F** data using GraphPad Prism and Tukey’s Multiple Comparison tests were used to evaluate differences compared to no treatment or empty (no receptor) membrane conditions Error bars are the mean ± S.D. ns, not significant, * p < 0.05, ** p < 0.01, *** p < 0.001, **** p < 0.0001.

We next examined TA peptidomimetic activation of the GPR97 holoreceptor and GPR97-CTFΔTA6 in HEK293T cells. A small screen of different stalk length peptides was performed and P7 (*i.e.* the first 7 residues of the GPR97 TA stalk) was identified as the most efficacious activator, whereas P18 had negligible activity, and ended up serving as a negative control in subsequent studies (Supplemental Figure 1). P7, but not P18 provided concentration-dependent activation of GPR97 and GPR97-CTFΔTA6 in the CRE-LUC and SRE-LUC assays (Figure 3C-3D). These results were corroborated using the GPR97-CTFΔTA6 G13 and Gs reconstitution assays, where P7 provided significant activation of the receptor and P18 had no activity (Figure 3E-3F).

### Comparison of tethered-agonist-bound and steroid-bound Adhesion GPCR structures

An alternative mode of adhesion GPCR activation to tethered agonist stimulation was proposed; steroids act with exquisite specificity as agonists for select adhesion GPCRs (12–15, 25). As a prelude to directly measuring steroid and tethered agonist GPR97 activation, AGPCR / steroid complex structures were aligned and superimposed (Figure 4C and Supplemental Table 1). We first examined the structures of GPR97 bound to beclomethasone and cortisol, as the corticosteroids were purported to activate GPR97 with high picomolar potency inhibition of cAMP production (12). Neither steroid stabilized a fully active state of GPR97 when compared to the active states of adhesion GPCRs bound to tethered peptide agonists (Figure 4A and 4B) (7, 11, 14). The GPR97 / steroid structures have rigid TM6 and TM7 α-helices and tethered agonist-bound adhesion GPCR structures have kinked TM6 and TM7 α-helices to permit access at the intracellular opening of the 7TM barrels for strong G protein engagement. Additional steroid-bound adhesion GPCR structures display kinked TM6 and TM7 α-helices (13, 14). However, an overlay of the available steroid-bound adhesion GPCRs demonstrates great positional variance of the steroids with no common binding pocket and non-specific hydrophobic interactions that mediate steroid binding to the receptors (Figure 4C). Despite the high chemical similarity of beclomethasone (BCM) and cortisol, the two steroids were bound to GPR97 in ∼180° horizontally-flipped orientations (Figure 4C) (12). DHEA was bound to GPR64 in two major orientations that differ in binding pocket depth and were both ∼180° vertically-flipped in comparison to BCM/Cortisol in complexes with GPR97 (Figure 4C) (12, 13). 5α-dihydrotestosterone (5α-DHT) and methenolone were presented to be exclusive binders of GPR133, but displayed three markedly distinct binding poses, vertical, ∼60° tilted from vertical and nearly horizontal and rotated (Figure 4C) (14). The variable poses of the different steroids are in stark contrast to the highly conserved conformation and binding interactions that TAs have in adhesion GPCR orthosteric sites (7, 8, 11, 31).

**Figure 4:**
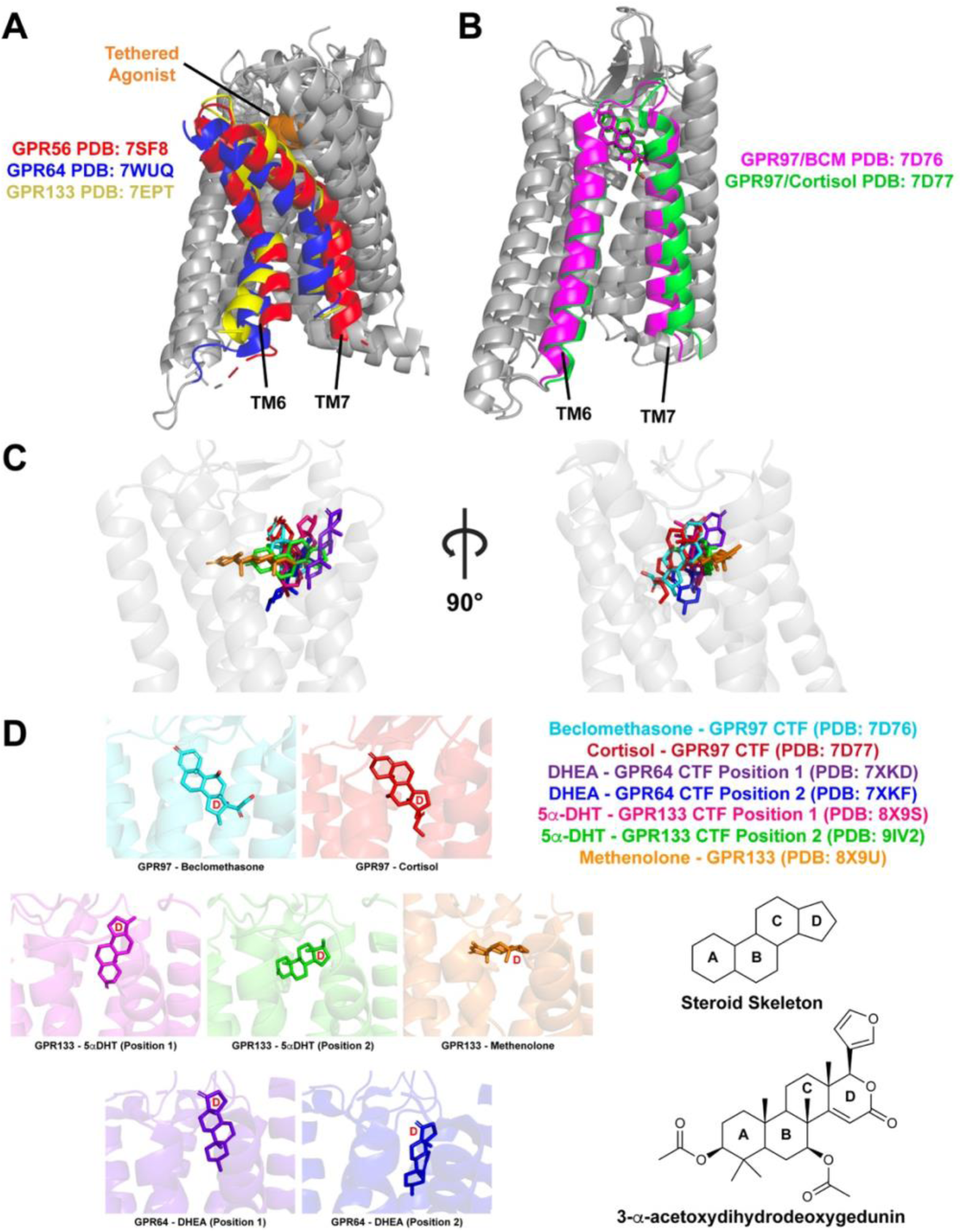
Comparison of tethered agonist-bound and steroid-bound adhesion GPCR structures. (**A**) Overlay of GPR56/ADGRG1, GPR64/ADGRG2, and GPR133/ADGRD1 active-state structures depicting intramolecular binding of the tethered agonists (orange) and the kinked TM6 and TM7 α-helices. (**B**) Overlay of GPR97/ADGRG3 partially active-state structures depicting beclomethasone (pink) and cortisol (green) binding and the rigid-body TM6 and TM7 α-helices. (**C**) Overlay of adhesion GPCR structures depicting the variable conformations of bound steroids, modeled using beclomethasone-bound GPR97 (PDB: 7D76). (**D**) Individual structural views of steroid orientations bound hydrophobically to adhesion GPCRs. The D-ring of the steroid skeleton is denoted to orient the modes of steroid binding. The chemical structure of the partial agonist, 3-α-DOG is depicted for comparison.

To further assess the positional variance of steroid binding to GPR97 we performed unbiased docking of BCM and additional steroids including 17-α-hydroxypregnenolone that was described to have no ability to activate GPR97 (12, 32). Multiple positions for each steroid were predicted proximal to the GPR97 orthosteric site with similarly favorable docking scores (kcal / mol) (Supplemental Figure 2A). The top 9 highest scoring poses of BCM and 17α-hydroxypregnenolone docked to GPR97 were compared to portray the variability (Supplemental Figure 2B). In sum, the high variability of steroid binding poses for multiple adhesion GPCRs suggests that binding may have been driven by incubation of high concentrations of hydrophobic steroids (*e.g.* 10 µM) with the receptors during the preparation phases prior to structure determination (12–14). This calls steroid-mediated adhesion GPCR signaling into question and warrants a direct comparison of steroid and tethered agonist efficacies.

### Direct comparison of tethered agonist, steroid, and the ADGRG partial agonist, 3-α-DOG activation of GPR97/ADGRG3

To compare steroid and tethered agonist activation of GPR97 we selected a mini-panel of steroids that were purported to have potent and exclusive action towards individual ADGRG family receptors including GPR97/ADGRG3, GPR56/ADGRG1, GPR64/ADGRG2, and GPR126/ADGRG6 (12, 13, 15, 33). We also compared the action of 3-α-DOG, a compound obtained by high throughput screening for GPR56 activators and found to be a partial agonist for ADGRG1/GPR56 and ADGRG5/GPR114, but not ADGRF family members (29, 30). CRE-LUC or SRE-LUC assays were first calibrated to the maximal efficacies obtained using tethered agonist-activated GPR97-CTF or GPR56-CTF constructs (Figure 5, green lines). GPR97 holoreceptor, GPR97-CTFΔTA6, or GPR56-CTFΔTA3 were then tested for activation with concentration series of the indicated steroids and 3-α-DOG (Figure 5A – 5E). 3-α-DOG exhibited partial agonism for GPR97 and GPR56 with potencies in the mid-micromolar range. At high concentrations, progesterone and DHEA exhibited weak activation of GPR97 (Figure 5A, 5B, 5D). None of the steroids purported to activate GPR97 including BCM, dexamethasone, cortisone, or hydrocortisone exhibited any activities in our assays (12). None of the steroids activated GPR56-CTFΔTA3 despite 17-α-hydroxypregnenolone being described as a potent and exclusive activator of the receptor (Figure 5E) (15). To validate these findings, we measured the kinetics of GPR97-CTFΔTA6 stimulated G13 GTPγS binding in the presence of 25 µM steroids or 3-α-DOG (Figure 5F). The steroids provided no activation in this orthogonal assay, whereas 3-α-DOG exhibited its established partial agonism.

**Figure 5:**
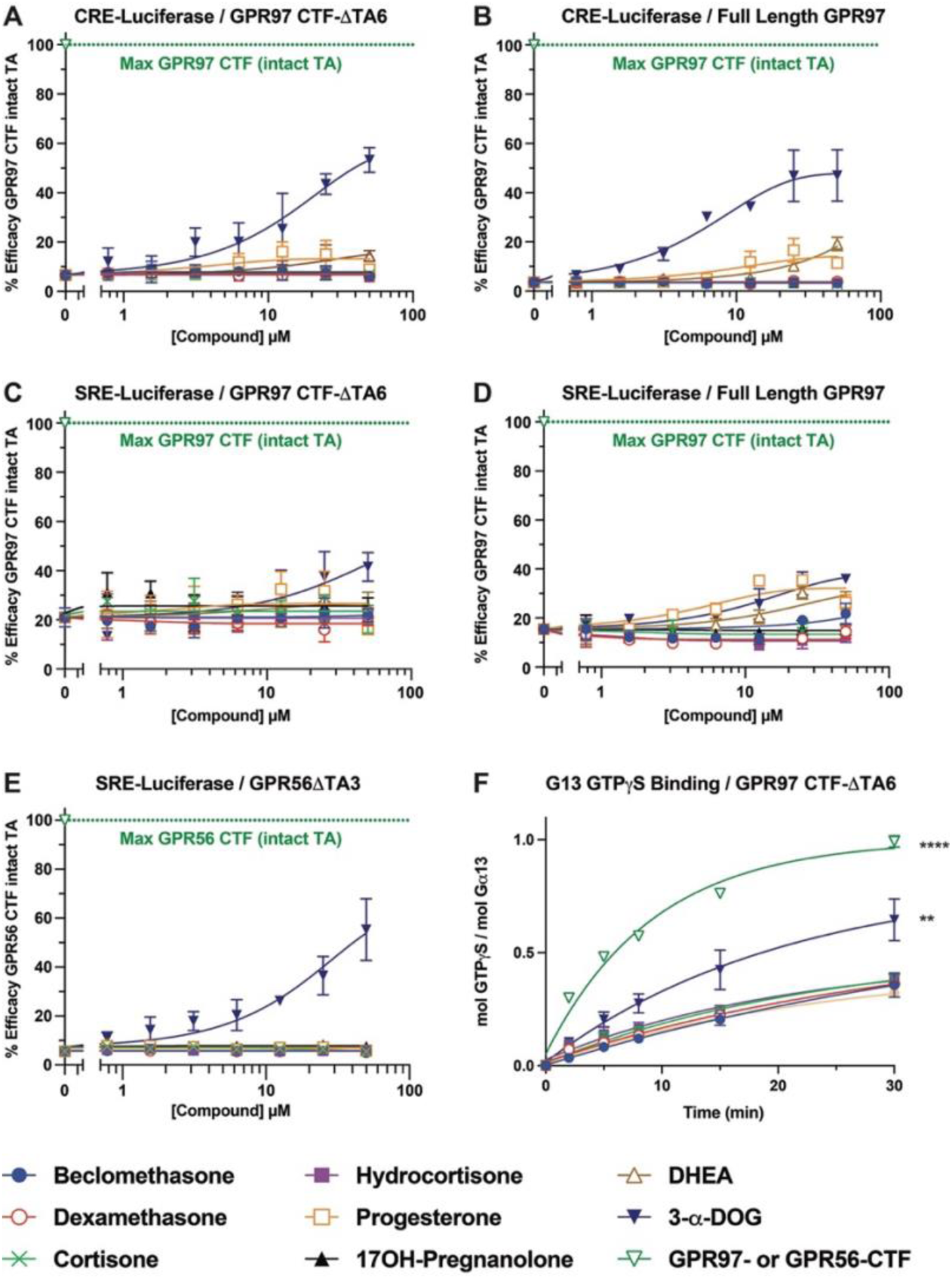
3-α-DOG is a partial agonist for GPR97 while steroids exhibit non-specific minimal activation of GPR97 and GPR56. HEK293 cell CRE-Luciferase activity measurements of full tethered agonist-stimulated GPR97-CTF (green line) compared to **(A)** GPR97-CTFΔTA6 and (**B**) full length GPR97 stimulated by 3-α-DOG or the indicated steroids. HEK293 cell SRE-Luciferase activity measurements of full tethered agonist-stimulated GPR97-CTF (green line) compared to **(C)** GPR97-CTFΔTA6 and (**D**) full length GPR97-stimulated by 3-α-DOG or the indicated steroids. **(E)** HEK293 cell SRE-Luciferase activity measurements of full tethered agonist-stimulated GPR56-CTF (green line) compared to GPR56-CTFΔTA3-stimulated by 3-α-DOG or the indicated steroids. (**F**) G13 GTPγS binding kinetics stimulated by full tethered agonist-stimulated GPR97-CTF (green line) compared to GPR97-CTFΔTA6 stimulated by 3-α-DOG or the indicated steroids. All data points depict the mean of biological triplicates. Error bars show the S.D. One-way ANOVAs were used for statistical analyses. ns, not significant, * p < 0.05, ** p < 0.01, *** p < 0.001, **** p < 0.0001.

### GPR97 activation induces human and mouse neutrophil polarization

GPR97 was found in neutrophils by RNA expression analysis and receptor immunoblotting and flow cytometry analysis (19, 34). We verified GPR97 expression by performing a comparative Taqman adhesion GPCR gene expression array of mRNA harvested from human platelets and human polymorphonuclear neutrophils (hPMNs). GPR56 mRNA was the most abundant of adhesion GPCRs in platelets, coinciding with its role in sensing collagen during hemostasis (2). GPR97 mRNA ranked fourth among five adhesion GPCRs detected in hPMNs, behind the leukocyte cell surface antigens, EMR2 and EMR3, and Latrophilin-3 (Supp Figure 3A). GPR97 protein was detected by western blotting in hPMN and mouse bone marrow neutrophil (mBMN) membrane homogenates (Supp Figure 3B). GPR97 was previously linked to Go signaling (12, 19) and activation of G12/13 effector enzymes (35); however its function in neutrophils, particularly in light of our findings that it couples strongly to G13 and Gs, remains unexplored. One of the first steps neutrophils undergo during capture by the vascular endothelium in preparation for extravasation is to acquire a polarized shape through signaling to the actin cytoskeleton (36). We assessed GPR97 agonist-dependent mBMN and hPMN polarization by treating freshly isolated neutrophils with the active and inactive GPR97 TA peptidomimetics P7 and P18, 3-α-DOG, or BCM – the potent neutrophil chemoattractant N-formylmethionine-leucyl-phenylalanine (fMLF) was used as a positive control (Figure 6) (37). Neutrophils were fixed and stained with DAPI and phalloidin to visualize polymerized actin. The two active GPR97 agonists, P7 and 3-α-DOG, but not inactive P18 or BCM, strongly induced neutrophil polarization in a manner that was comparable to the response induced by fMLF when fields of neutrophils were inspected qualitatively (Figure 6A and 6D). Shape change and polarization of mBMNs (Figure 6B-6C and Supp Figure 4) and hPMNs (Figure 6E-6F and Supp Figure 5) were quantified by scoring changes in cell area and cell circularity. P7, 3-α-DOG, and fMLF significantly altered these parameters, whereas P18– and BCM-treated neutrophils remained small and circular, much like vehicle-treated cells. We observed that 3-α-DOG appeared to induce more neutrophil filipodia protrusions, compared to the two stronger agonists P7 and fMLF (see Figure 6A and D). We speculate that this may be due to an off-target effect of 3-α-DOG, which is known to activate all ADGRG adhesion GPCRs tested to date (Figure 5) (29, 30).

**Figure 6:**
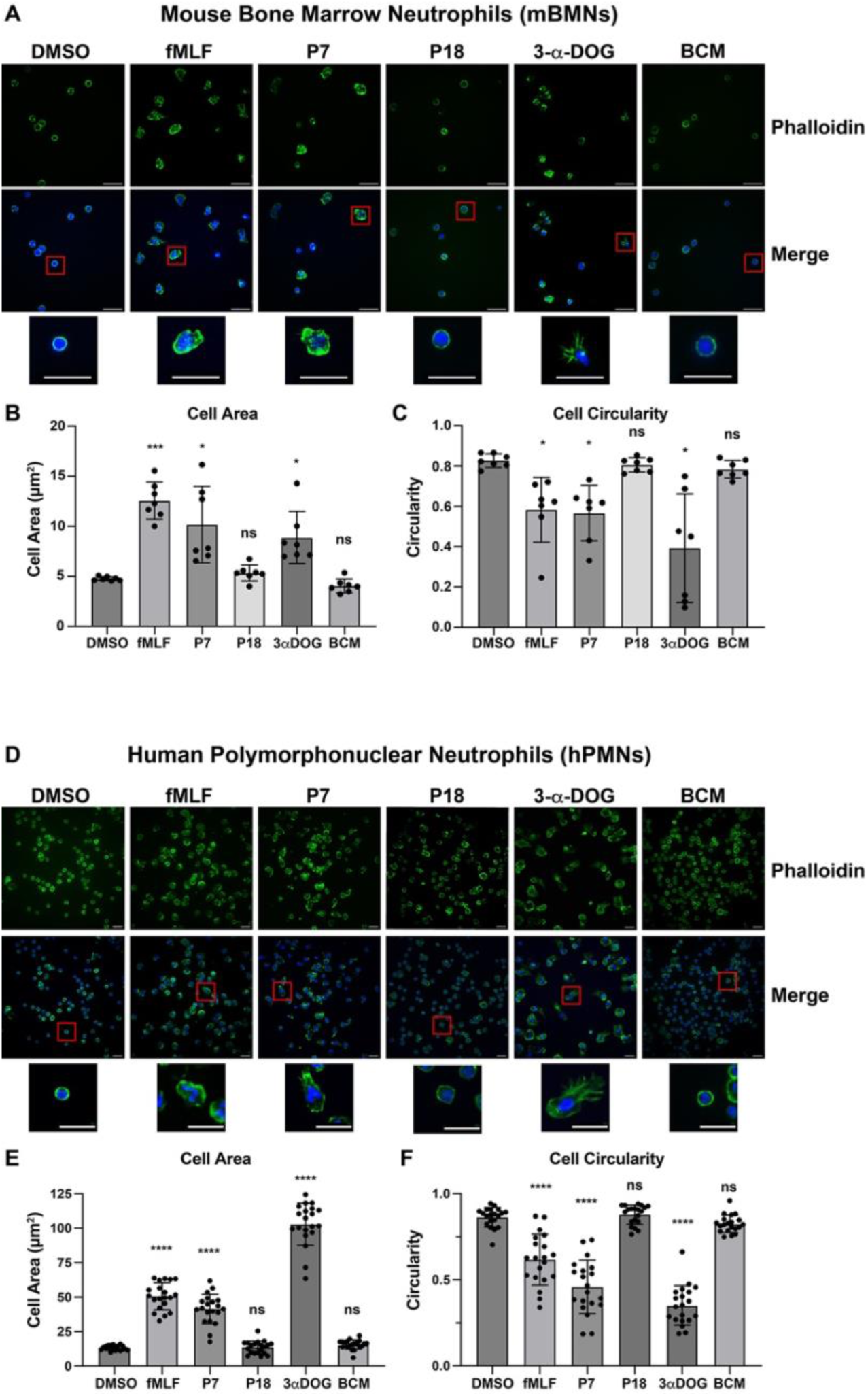
GPR97 activation induces neutrophil polarization. Mouse bone marrow neutrophils (**A-C**) and human PMNs (**D-F**) were treated with N-formylmethionine-leucyl-phenylalanine (fMLF), the active (P7) and mock (P18) GPR97 TA peptidomimetics, 3-α-DOG, beclomethasone (BCM) or vehicle control (DMSO), stained with phalloidin and DAPI and imaged by confocal microscopy. Scale bars are 20 µm. Image quantification of cell surface area (**B and E**) and circularity (**C and F**) for each condition. Error bars are the mean ± S.D. hPMNs n=20. mBMNs n=7. One-way ANOVAs were used for statistical analysis in comparison to DMSO. ns, not significant, * p < 0.05, ** p < 0.01, *** p < 0.001, **** p < 0.0001.

### GPR97 activation induces human and mouse neutrophils chemotaxis

Neutrophil activation, polarization, and chemotaxis are mediated by a complex orchestration of multiple G protein signaling pathways and transient adhesive interactions with the endothelium and smooth muscle cells that comprise vascular vessel walls as well as with various interstitial cells (36, 38). We examined GPR97-agonist induced mBMN and hPMN chemotaxis using two approaches, trans-well migration and the agarose spot assays (Figure 7). Fresh isolated neutrophils were placed in the upper chambers of trans-well devices and the cells that migrated through the porous membranes were quantified after 2 hrs. We found that both human and mouse neutrophils showed a strong response towards P7, comparable to the response towards fMLF, and a somewhat lesser response towards 3-α-DOG (Figure 7A-7B). No increase in migration above vehicle control was observed for P18 or BCM.

**Figure 7:**
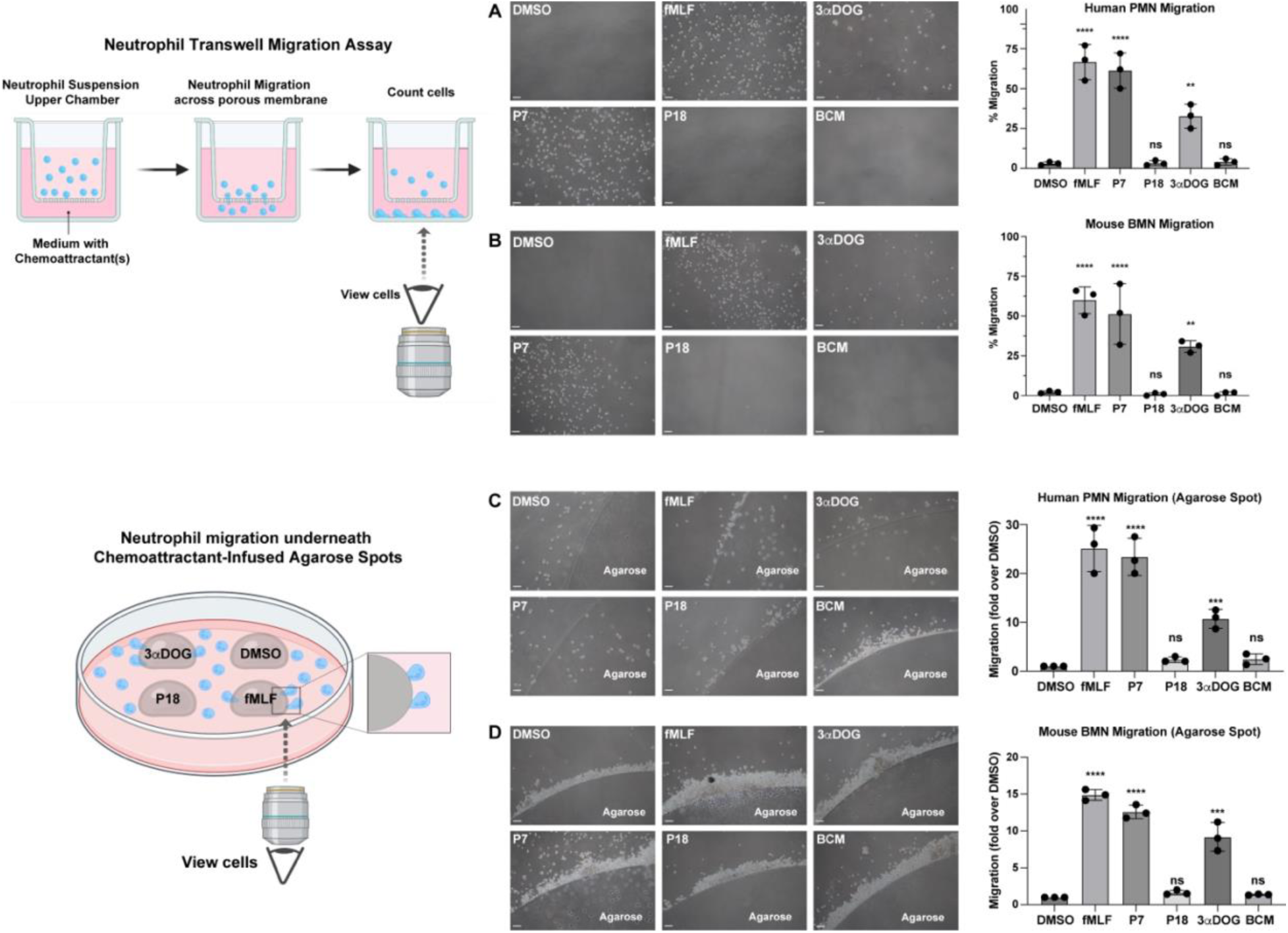
GPR97 activation induces neutrophil migration. (**A**) hPMN and (**B**) mBMN were induced to migrate across a Transwell chamber membrane toward fMLF, the active (P7) or inactive (P19) GPR97 TA peptidomimetics, 3-α-DOG, BCM, or vehicle control (DMSO). The micrographs depict representative fields of neutrophils that migrated to the lower chambers containing the agonists. Data are the percentage of cells migrated and are biological triplicates. Scale bars are 50 µM. (**C**) hPMN and (**D**) mBMN migration in the chemotactic agarose spot assay in response to fMLF, the active (P7) or inactive (P19) GPR97 TA peptidomimetics, 3-α-DOG, BCM, or the vehicle control (DMSO). Scale bars are 50 µM. Data are presented as a ratio of total cells migrated under the agarose spot over DMSO cells migrated. Error bars are the mean ± S.D. One-way ANOVAs were used for statistical analyses in comparison to DMSO. ns, not significant, * p < 0.05, ** p < 0.01, *** p < 0.001, **** p < 0.0001.

Similar results were obtained using the agarose spot assay in which chemoattractants were infused into agarose and the gelled agarose spots were placed into a culture dish with neutrophils. In this assay, neutrophils build up on the outside of the agarose spots under all conditions, but when the neutrophils sense the chemoattractant, they respond by crawling underneath the agarose spot and migrate directionally towards its center, which was scored by counting the total cells under the agarose spots (Figure 7) (39). Migration of mBMNs (Figure 7B-7D) and hPMNs (Figure 7A-7C) was prevalent in the fMLF and P7 conditions but not in the DMSO or P18 conditions. As for the transwell assay, 3-α-DOG induced partial migration of mBMNs and hPMNs. These results show that neutrophil activation induced by two GPR97 agonists characterized here (P7 and 3-α-DOG) evokes the G protein signaling pathways that induce neutrophil shape change, polarization and chemotaxis. Steroids that did not activate GPR97 signaling in our hands were also unable to promote neutrophil chemotaxis.

## DISCUSSION

Understanding of the GPR97 G protein coupling profile has been mixed. Structural and signaling studies indicated that GPR97 binds to Go and inhibits adenylyl cyclase (AC), whereas another report suggested that basal activity of the holoreceptor stimulated AC through Gs, while an expressed version of the CTF activated Gi (12, 19, 25). A third study showed that GPR97 activated effector enzymes that are downstream of G12/13 (35). We took a reductionist approach to comparatively and directly measure GPR97 coupling to representative members of all four G protein families and demonstrated that GPR97 strongly activated G13 and Gs, provided appreciable activation of Gi_1_, and did not couple to Gq (Figure 1). These findings were bolstered using orthogonal assays in heterologous cells; GPR97 strongly stimulated the SRE-LUC and CRE-LUC reporters which are activated downstream of G13 and Gs, respectively (Figure 3).

We found that GPR97 is a canonical adhesion GPCR as its self-cleavage and the dissociation of its NTF and CTF are requisite for activation by its tethered-peptide-agonist, which has now been verified for all members of the ‘G’ subfamily of adhesion GPCRs that have been tested (7, 8, 10, 29). This includes GPR56(G1) and GPR114(G5), the closest homologs of GPR97(G3), which appear to be the products of genetic duplication as the genes are aligned in tandem on human chromosome 16. This hints at a potential point of evolutionary pressure for the gene duplication events; each gene product may regulate a specialized function on select differentiated cells of the hematopoietic stem cell (HSC) lineage. GPR114 and GPR97 are among the simplest adhesion GPCRs with minimal GAIN domains as NTFs and have limited expression patterns, primarily in eosinophils and leukocytes/neutrophils, respectively (19, 29, 40). GPR56 is distributed broadly and its NTF has the added complexity of an N-terminal <u>p</u>entraxin/laminin/neurexin/sex-hormone-binding-globulin-like domain (PLL) that binds to collagen (3, 41). This coincides with its high abundance on the surface of platelets, where it assists platelets in collagen-dependent activation during hemostasis as a blood flow-dependent shear force transducer (2, 42). The white cell lineages that express GPR97 and GPR114 are not regulated like platelets by the collagen mesh that surrounds blood vessels, rather these cells interact with the vascular endothelium prior to undergoing TEM.

Future investigations will explore the possibility that GPR97 and GPR114 (29) mediate shear force-dependent endothelial capture and polarization of neutrophils during TEM by interacting with putative ligands presented by the endothelium or by additional cells that comprise vessel walls. For example, as a neutrophil progresses through TEM, the GPR97 NTF may become attached to ligands presented by various vessel wall cells, and the processes of rolling along the endothelium or continued migratory movement could provide shear force to dissociate the GPR97 NTF and allow for rapid TA-mediated receptor activation. This would evoke neutrophil G13, Gs, and Gi signaling pathways to orchestrate neutrophil actin cytoskeleton-dependent polarization, migration, and regulation of cAMP levels that can be pro-or anti-migratory depending on context (22, 43, 44).

To address this, we conducted a primary investigation of GPR97 involvement in neutrophil polarization and chemotaxis by stimulating human and mouse neutrophils with GPR97 agonists. Stimulation with the established agonist fMLF, evoked neutrophil polarization and migration in two independent assays that was comparable to that induced by P7 and to a lesser degree, 3-α-DOG (Figure 6 and 7). The fMLF receptor couples to Gi and its roles in Gαi and Gβγ-dependent neutrophil polarization and migration are well established (45). High cAMP levels are generally considered inhibitory to neutrophil migration, however, there are instances where small pulses of Gs-dependent cAMP production help to initiate migratory processes through actin cytoskeleton contraction (43, 46). As GPR97 is coupled to G13, Gs and Gi/o, its signaling role in neutrophil polarization and migration is complex. Our results clearly show that GPR97 stimulus overall is pro-migratory. However, it is not clear whether GPR97 has a direct role in chemotaxis or works as a chemokinetic receptor. It seems more plausible that GPR97 may work in concert with known directionally-sensing chemotactic receptors. Future work with *Gpr97* deletion models, use of G protein pathway specific inhibitors, and assays that measure neutrophil migration *in vivo* or in the presence of endothelial cells will help to pinpoint the exact role of GPR97 in neutrophil TEM.

The corticosteroid beclomethasone was purported to activate GPR97 but had no effect in our assays of neutrophil polarization or migration when compared to *bona fide* agonists of the receptor (Figure 6 and 7) (12, 25). A new controversy has sought to position adhesion GPCRs as targets for the long sought, non-genomic actions of steroids (12–15, 33). Despite being chemically similar, individual steroids were purported to activate select adhesion GPCRs with high potency (*e.g.* pM to nM) and exclusive specificity (12). In addition to showing no effects of BCM in the neutrophil functional assays, we used cell-based and *in vitro* biochemical assays to demonstrate that mid-µM concentrations of BCM and related steroids provided negligible activation of GPR97 in comparison to its tethered agonist or 3-α-DOG. In fact, steroids that were purported to exclusively activate other adhesion GPCRs besides GPR97 (*e.g.* progesterone and DHEA) marginally improved GPR97 activation efficacy above that of BCM / vehicle control levels, but were far below the efficacy imparted by the full tethered agonist (Figure 5) (12, 25).

We compared solved structures of steroid-bound GPR97, GPR64, and GPR133 to illustrate the high variances of the steroid binding poses (12–14). The D-ring of each steroid was denoted as a visualization point and it was illustrated that the D-ring of DHEA was closest to the extracellular face of the GPR64 7TM bundle, while BCM and cortisol were rotated 180° in relation to the DHEA pose and their D rings were oriented towards the core of the GPR97 7TM bundle (Figure 4). Moreover, BCM and cortisol were rotated 180° horizontally in relation to each other within the GPR97 7TM bundle. 5α-DHT and methenolone were depicted within the GPR133/ADGRD1 7TM bundle in three orientations vertical, angled, and horizontal (14). These dramatic variances in the positions of the steroids bound to adhesion GPCRs suggests that there are no defining characteristics of the steroid chemical structures or the receptors that account for the claimed specificity and exclusivity of adhesion GPCR target binding. Adhesion GPCR orthosteric sites evolved to accommodate the highly hydrophobic seven amino acid tethered peptide agonists. It may be that steroids were experimentally induced to occupy these large hydrophobic cavities within adhesion GPCRs in situations where the peptide agonist is absent either through molecular engineering or when it is sequestered within the GAIN domain. Additional testing of steroid action on adhesion GPCRs by multiple laboratories may help to sort out the disparate observations.

## MATERIALS AND METHODS

### Molecular Cloning

Human GPR97 cDNA (UniProt Accession Number Q86Y34) was used as a template for PCR-based subcloning of full length, truncated, and mutagenized cDNAs into pFastBac-1 or pcDNA3.1 expression vectors as indicated. Baculoviruses were generated following the Bac-to-Bac system manufacturer’s instructions (Invitrogen).

### Sf9 culture, baculovirus and recombinant AGPCR membrane production

*Spodoptera frugiperda* 9 (*Sf*9) cells were cultured in shake flasks at 27 °C in ESF921 medium (Expression Systems). Recombinant bacmids were produced using the Bac-to-Bac system (Invitrogen) and transfected into *Sf*9 cells using Fugene HD (Promega). Baculoviral supernatants were harvested five days after transfection. For viral amplification, *Sf*9 cells grown to 2.0-3.0 x 10^6^ cells/mL were infected with a 1/100 dilution of virus. Amplified viruses were harvested 72-96 h post-infection.

For AGPCR membrane homogenate preparation, *Sf*9 cells growing at 2.0-3.0 x 10^6^ cells/ml were infected with 1/50^th^ volume of amplified baculovirus and harvested 48 h post-infection. *Sf*9 AGPCR cell pellets were resuspended in lysis buffer (20 mM HEPES pH 7.4, 1 mM EDTA, 1 mM EGTA, and protease inhibitor cocktail (23 mg/mL phenylmethylsulfonyl fluoride, 21 mg/mL Nα-*p*-tosyl-L-lysine-chloromethyl ketone, 21 mg/mL L-1-*p*-tosylamino-2-phenylethyl-chloro ketone, 3.3 mg/mL leupeptin, and 3.3 mg/mL lima bean trypsin inhibitor). Cells were lysed using a nitrogen cavitation device (Parr Industries). The lysates were centrifuged at 600 g for 10 min and the resultant supernatant was centrifuged at 100,000 g for 40 min at 4 °C. The membrane pellet was split into two samples and Dounce homogenized at 4°C into lysis buffer or lysis buffer containing 7M urea for NTF solubilization. The samples were then centrifuged at 100,000 g for 40 min at 4 °C. The membrane pellet washed of urea by Dounce homogenization into 20 mM HEPES pH 7.4, 1 mM EGTA, recentrifuged and Dounce homogenized into storage buffer, 20 mM HEPES pH 7.4, 1 mM EGTA 11% w/v sucrose. The concentration of total protein in the membrane homogenates was determined by Bradford assay.

### SDS-PAGE and Immunoblotting

AGPCR cell membranes (50 µg) were treated ± 500 units of PNGaseF (New England Biolabs Cat# P0704S) for 1 hr at 30 °C. Membranes were then mixed with reducing SDS-sample buffer and resolved by 12% SDS-PAGE followed by transfer to PVDF membranes for immunoblotting. Samples were not heated prior to gel loading. Chemiluminescent western blotting was used to detect wild type and mutant GPR97. GPR97 polyclonal antibody (Cat# PA5-116252) was used to detect the human N-terminal fragment of GPR97 (1:2000). Invitrogen GPR97 Polyclonal Antibody (Cat# PA3-049) was used to detect the human and mouse C-terminal fragment (1:2000). The secondary antibodies used were anti-Rabbit IgG peroxidase-linked antibody from donkey (1:5000, Fisher Scientific) and anti-Mouse IgG peroxidase-linked antibody from donkey (1:5000, Fisher Scientific). Blot images were acquired using an iBright FL1000 system (Invitrogen).

### Luciferase Reporter Assays

Low-passage HEK293T cells (ATCC CRL-3216) were plated at 9.0 x 10^6^ cells per 15 cm dish in DMEM + 10% (v/v) FBS 24 hours prior to polyethylenimine (PEI)-mediated transfection with 3 µg of AGPCR in pcDNA3.1, 8.22 µg SRE luciferase reporter (pGL4.33) (Promega) or CRE-luciferase reporter (CRE-luc2p Promega) and 82.5 ng of phRLuc-N1 (PerkinElmer). Each transfection was balanced to 26.5 µg total plasmid DNA with additional pcDNA3.1. The transfected cells were lifted 6 h later with 0.05% w/v trypsin in Puck’s Saline G containing 1 mM EGTA (PSG-EGTA), pooled, frozen and thawed as needed to seed white, sterile 96-well, TC-treated plates (BrandTech BRAND) at 40,000 cells per well in serum-free DMEM for SRE-luciferase assays. For CRE-luciferase assays, the serum-containing medium was aspirated at 16 h and replaced with 100 µL of serum-free DMEM for compound or peptide dosing. 200 µM of steroids, 3-α−DOG, or GPR97 synthetic tethered agonist peptidomimetics were serial diluted in medium in a separate master plate. 100 µL of 2X concentrations were then added to the assay plate and incubated with the cells for an additional 4 h. The final DMSO content did not exceed 1% v/v. The plates were centrifuged for 5 min at 500 g and 150 µL of medium was removed. Cells were lysed by addition of 50 µL Steady-Glo reagent (Promega) and luminescence was measured using a TriStar2 plate reader (Berthold Technologies) (10). 100 µL *Renilla* luciferase (RLuc) buffer (1.1 M NaCl, 2.2 mM EDTA, 0.44 mg/mL bovine serum albumin, 0.22 M KH_2_PO_4_ pH 5.1) containing 3 µM coelenterazine h was added to each well prior to measuring the RLuc signal (Dryer et all 2000). Data were presented as the ratio of Firefly Luciferase (Fluc) units to Renilla Luciferase (RLuc) units.

### AGPCR / G protein Signaling Reconstitution Assays

Mock (5 µg / assay point) or urea-treated (equiv. vol.) AGPCR or GPCR membranes were reconstituted with purified Gαs, i_1_, q, or 13 (100 nM) and Gβ_1_Gγ_2_ (500 nM) in pre-incubation buffer containing 50 mM HEPES, pH 7.4, 1 mM DTT, 1 mM EDTA, 20 µM GDP, 3 µg/ml BSA. 25 µM of steroid, 3-α-DOG, or synthetic TA peptidomimetic were added as indicated. The G protein-reconstituted AGPCR membranes were incubated on ice and then at 25 °C, 2 min prior to initiation of triplicate reactions by addition of an equal volume of buffer containing 50 mM HEPES, pH 7.4, 1 mM DTT, 1 mM EDTA, 20 µM GDP, 3 µg/ml BSA, 10 mM MgCl_2_, 50 mM NaCl, 2 µM [^35^S]-GTPγS (20,000 cpm/pmol). Samples were removed as technical triplicates at times 2, 5, 8, 15 and 30 min and quenched in buffer containing 20 mM Tris (pH 7.7), 100 mM NaCl, 10 mM MgCl_2_, 1 mM GTP, and 0.08% (m/v) deionized polyoxyethylene 10 lauryl ether C12E10. The samples were then filtered onto Whatman GF/C glass microfiber filters (Cytiva) using a Brandel harvester, dried and subjected to liquid scintillation counting.

### Computational simulated docking of steroids to GPR97

Docking was conducted using the open-source MolModa 1.0.1 (Durrant Lab) (32). The resolved structure of the beclomethasone-bound GPR97 (PDB: 7D76) was used as the receptor base while various steroid compounds were docked within a 20Å^3^ grid centered around the upper half of the 7TM barrel proximal to the orthosteric site. Multiple docked positions were outputted with structures and docking scores in units of kcal/mol were obtained. Poses of BCM and 17-α-hydroxypregnenolone used for portrayal were selected based on the top scores generated.

### PyMol Structural Alignments

Resolved AGPCR structures were imported into PyMol from the Protein Data Bank (PDB) based on their PDB identification code. Structures were aligned to the GPR97-BCM structure (7D76) using the align function within PyMol. This function uses sequence information to align the structures followed by matrix calculations to minimize the Root Mean Square Deviation (RMSD). The maximum number of outlier rejection cycles was set to 5.0. The outlier rejection cutoff in RMSD was set to 2.0 Å.

### Mouse Bone Marrow Neutrophil Isolation

Femurs and tibias were dissected from deceased 8-16-week-old mice. The bone marrow was flushed out with Roswell Park Memorial Institute (RPMI)-1640 medium supplemented with 10% v/v FBS and 2 mM EDTA using a syringe with a 27G needle. The diluted marrow was centrifuged at 400 g to pellet cells. Cells were resuspended in 0.2% w/v NaCl to lyse red blood cells followed by the addition of 1.6% w/v NaCl to achieve isotonicity. Bone marrow cells were centrifuged at 400 g for 7 min and resuspended in RPMI-1640 with 10% v/v FBS and 2 mM EDTA and overlayed onto a Histopaque density gradient using 3 mL Histopaque 1119 (GE Healthcare BioSciences) and 3 mL Histopaque 1077 (Cytiva). This gradient was centrifuged at 400 g for 30 min. Neutrophils were collected from the middle layer and washed with DPBS before counting to determine cell viability using trypan blue dye exclusion.

### Polymorphonuclear Neutrophil Isolation

Deidentified, human blood was collected by the Department of Pharmacology Blood Core Facility into vacutubes containing lithium heparin. 25 mL of whole blood was mixed 1:1 with 3% w/v dextran from *Leuconostoc spp*. (Sigma-Aldrich Catalog No. 31392) solubilized in 0.9% w/v NaCl solution to enrich leukocytes by gravity sedimentation for 45 min. The leukocyte-rich layer was centrifuged in 1% w/v BSA-coated tubes at 400 g to further isolate leukocytes. The leukocytes were resuspended in 20 mL modified Hank’s Based Salt Solution (mHBSS) and underlaid beneath a 10 mL Ficoll-Paque (Cytivia) cushion for separation by centrifugation at 400 g for 30 min. The pelleted granulocytes and erythrocytes were resuspended in ammonium-chloride-potassium (ACK) Lysing Buffer for 5 min to lyse erythrocytes and recentrifuged. The PMN-rich pellet was resuspended in mHBSS, and cell viability was determined by trypan blue dye exclusion.

### Visualization of Neutrophil Polarization and Actin Polymerization

8-chamber cover glass chambers (Lab-Tek) were coated with 5 µg/mL fibronectin for 60 min at 37 °C. Isolated neutrophils were seeded at 2.5×10^5^ cells per chamber. 100 nM fMLF for hPMNs and 10 µM for mBMNs, 25 µM GPR97 tethered agonist peptidomimetics, 25 µM of beclomethasone and 3-α-DOG, or DMSO were added to the chambers and incubated for 10 min. Cells were perforated with 0.4% w/v saponin for 10 min fixed with 4% w/v paraformaldehyde (PFA) for 20 min and washed with Dulbecco’s Phosphate Buffer Solution (DPBS). The fixed cells were incubated with 1:500 Alexa Fluor 488-phalloidin (Invitrogen) and 1:2000 Hoesct 33258 dye (Invitrogen) for 30 min and washed DPBS. Images were acquired using a Leica DMi8 confocal microscope. Images processing and analyses were performed using ImageJ.

### Transwell Chamber Assays

Polyester inserts with 3.0 µm pore size (CellQUART) were placed into each well of 24-well non-tissue cultured treated plates. The wells were incubated with 5 µg/mL fibronectin solution for 60 min prior to experiments. Chemoattractant reagents (100 nM fMLF for hPMNs and 10 µM fMLF for mBMNs) or synthetic TA peptidomimetics (25 µM) in mHBSS were added into the bottom wells. Neutrophils (4.0 x 10^5^) were seeded into each insert and allowed to migrate for 60 min. Total migration to the bottom wells was determined by cell counting using a hemacytometer and imaging was performed using a Nikon Eclipse TS100 inverted scope with a 10X objective lens and a Canon EOS Rebel T7 with a 2.5X Universal DSLR to Microscope Adapter (Martin Microscope).

### Chemotactic agent Agarose Spot Assay

SeaKem LE Agarose (Lonza) was heated in 20 mL DPBS to make a 0.5% agarose molten gel. The gel was individually mixed with chemotactic agents, 25 µM TA peptidomimetics, 3-α-DOG, or beclomethasone, 100 nM fMLF for human PMNs, or 10 µM fMLF for mouse BMNs. Agent-infused 10 µL agarose drops were added to quadrants of a 20 mm glass bottom culture dish (Nest) and allowed to solidify at 4 °C. Isolated hPMNs (6.5 x 10^4^) and mBMNs (1.3 x 10^5^) in 2 mL of RPMI 1640 medium were added to the culture dishes. Cells were placed in a 37 °C, 5% CO_2_ tissue-culture incubator for 60-120 min. Brightfield imaging was performed using a Nikon TS100 Eclipse inverted scope with a 10X objective lens and a Canon EOS Rebel T7 with a 2.5X Universal DSLR to Microscope Adapter (Martin Microscope). The total number of cells migrated under each agarose spot were counted. Data was presented as a ratio over DMSO.

## STATISTICAL ANALYSIS

All data are presented as the mean ± S.D. of three or more independent experiments (biological replicates). GraphPad Prism was used for all statistical analyses. The threshold for significance was P < 0.05.

## Supporting information

Supplemental Section

## ACKNOWLEDGEMENTS

This work was funded by the Dr. Benedict and Diana Lucchesi Graduate Education Fellowship awarded to T.F.B., AHA grant 25POST1377764 to Y.F. and NIH grants R35GM149539 to G.G.T. and R35GM127303 to A.V.S.

