## Supplemental Section for "GPR97/ADGRG3 is activated by its tethered peptide agonist and not by steroids to induce neutrophil polarization and migration"

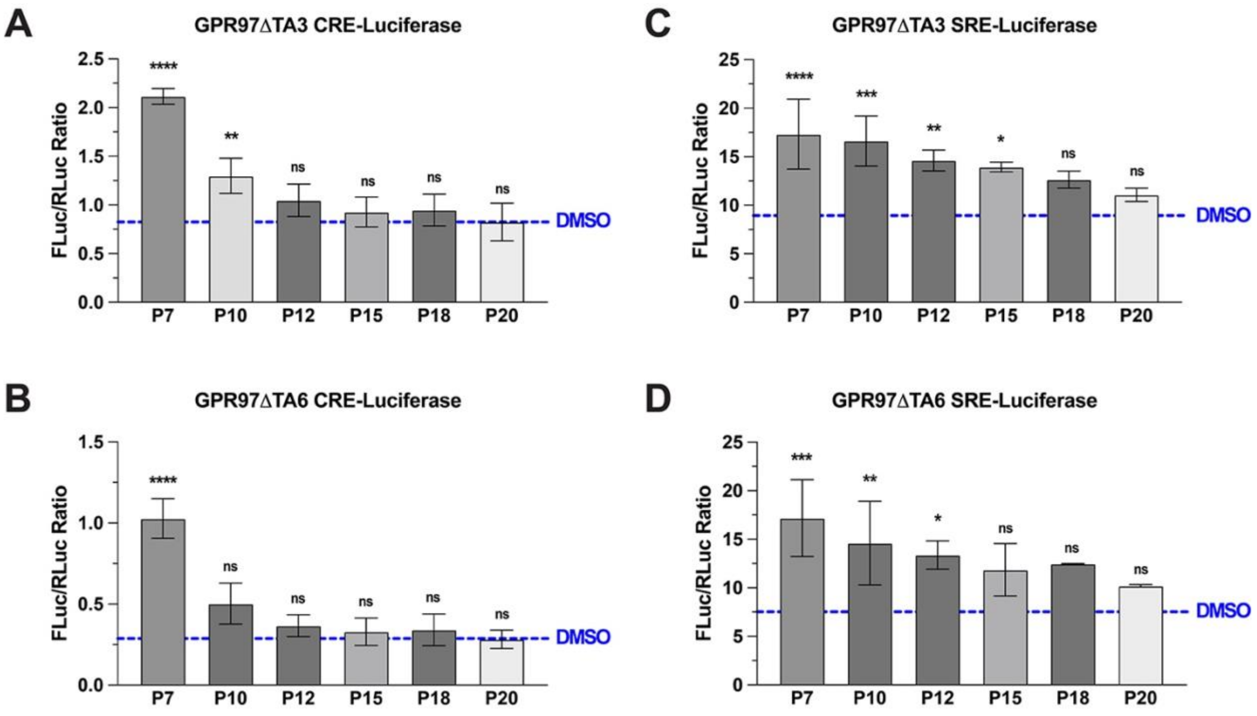


**Supplemental Figure 1:** GPR97 TA peptidomimetic-stimulated CRE-LUC activation for (**A**) GPR97-CTF-ΔTA3 and (**B**) GPR97-CTF-ΔTA6. GPR97 TA peptidomimetic stimulated SRE-LUC activation for (**C**) GPR97-CTF-ΔTA3 and (**D**) GPR97-CTF-ΔTA6. Dashed lines represent vehicle control (DMSO) as constitutive receptor activities. Data are presented as the ratios of the FLuc reporter signal to RLuc balancer signal and are biological triplicates Error bars are the mean ± S.D. One-way ANOVAs were used for statistical analyses. ns, not significant, * $p < 0.05$, ** $p < 0.01$, *** $p < 0.001$, **** $p < 0.0001$.

**A**

**Steroid : GPR97 Docking Scores (kcal / mol)**

| | Beclomethasone | Dexamethasone | Hydrocortisone | DHEA | 17-α-hydroxypregnenolone | Progesterone |
|---|---|---|---|---|---|---|
| **Model 1** | -10.352 | -10.587 | -9.511 | -9.759 | -9.886 | -10.141 |
| **Model 2** | -10.283 | -10.151 | -9.419 | -9.58 | -9.855 | -9.782 |
| **Model 3** | -10.073 | -10.043 | -9.414 | -9.37 | -9.693 | -9.764 |
| **Model 4** | -10.06 | -9.881 | -9.163 | -9.318 | -9.532 | -9.509 |
| **Model 5** | -10.03 | -9.718 | -9.127 | -9.28 | -9.527 | -9.425 |
| **Model 6** | -10.024 | -9.606 | -9.035 | -9.273 | -9.429 | -9.366 |
| **Model 7** | -9.976 | -9.378 | -9.031 | -9.202 | -9.393 | -9.343 |
| **Model 8** | -9.871 | -9.133 | -8.994 | -9.178 | -9.363 | -9.164 |
| **Model 9** | -9.649 | -9.12 | -8.951 | -9.098 | -9.326 | -9.015 |

**B**

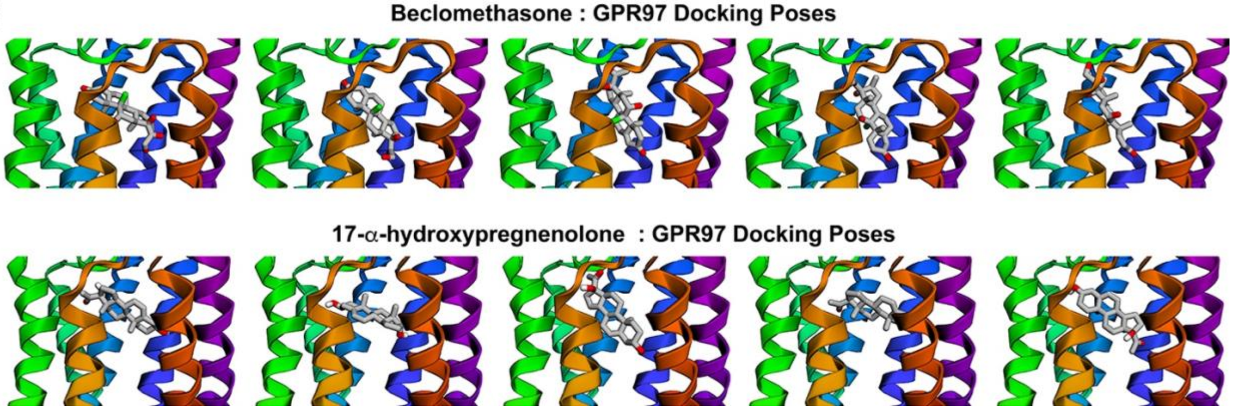


**Supplemental Figure 2: GPR97 steroid binding variances predicted by unbiased docking.** (**A)** MolModa open-source docking software scores (kcal/mol) for the top nine models of the indicated steroids docked to GPR97 (PDB: 7D76) (**B**) The top five docking poses of beclomethasone and 17-α-hydroxypregnenolone proximal to the GPR97 orthosteric site.

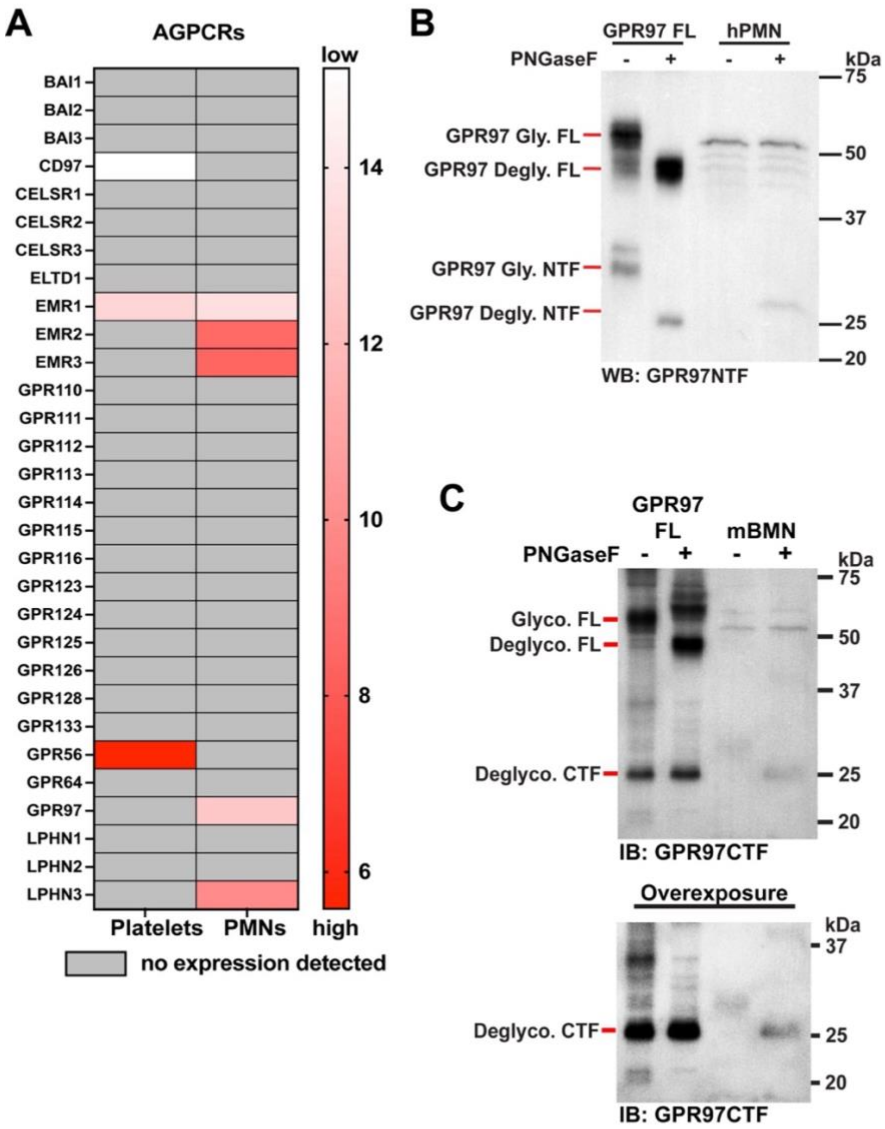


**Supplemental Figure 3: Adhesion GPCR mRNA expression in human platelets and neutrophils and GPR97 protein levels in human and mouse neutrophils.** (**A**) TaqMan GPCR array measurements of relative adhesion GPCR mRNA levels in human platelets and neutrophils. (**B**) Immunoblotting of overexpressed human GPR97 full length *Sf*9 membrane homogenates and human neutrophil membrane homogenates treated ± PNGaseF. We were unable to detect the glycosylated NTF band from neutrophils, which is consistent with our findings in failing to detect the endogenous GPR114 NTF in an eosinophil cell line (29). The endogenous deglycosylated GPR97 NTF band had a slightly higher MW than the overexpressed NTF. We do not know the reason for this difference other than it is not a shift caused by N-linked glycans. (**C**) GPR97WT overexpressed *Sf*9 membrane homogenates and mouse bone marrow neutrophil membrane homogenates were untreated or treated with PNGaseF and immunoblotted with a GPR97CTF specific antibody.

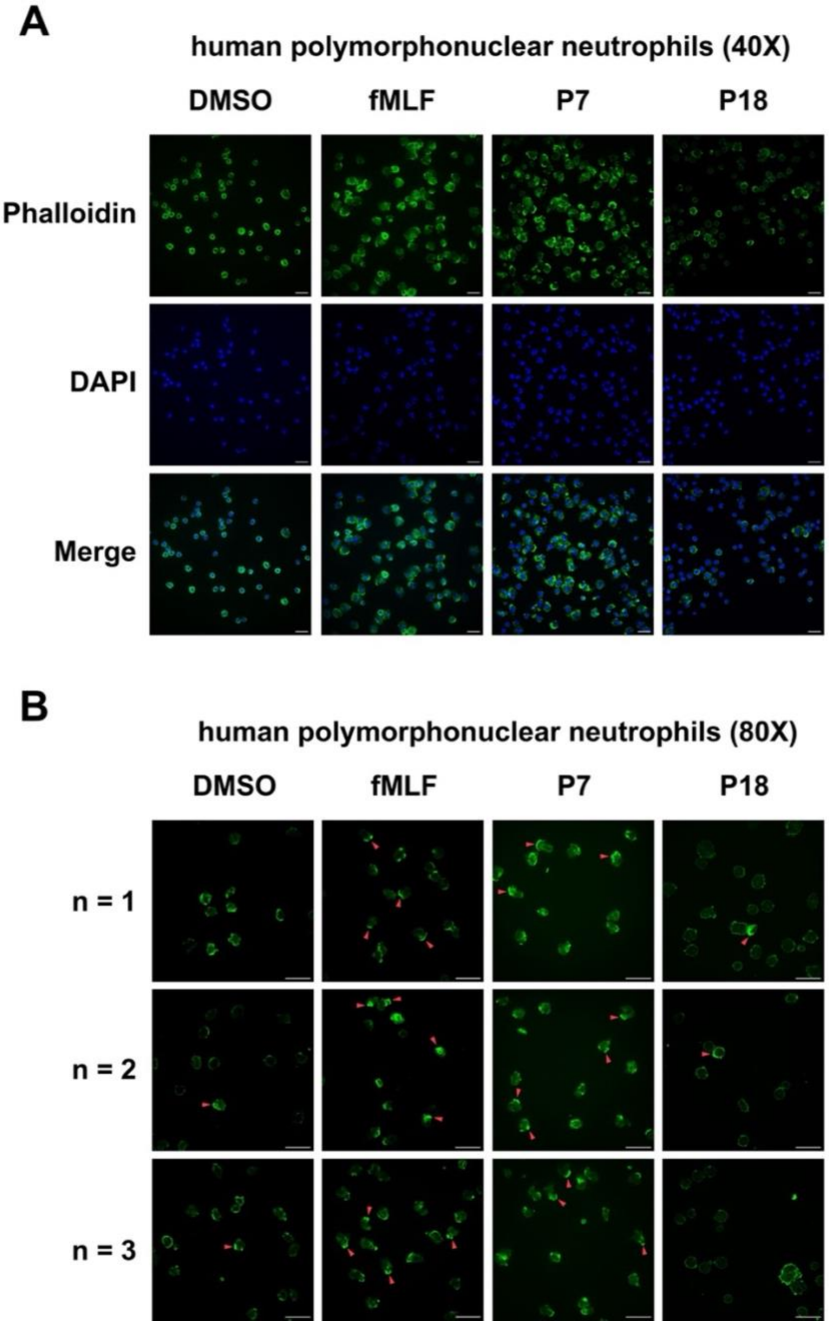


**Supplemental Figure 4: GPR97AP-dependent human neutrophil polarization.** hPMNs were stained with phalloidin and 4', 6'-diamidino-2-phenylindole (DAPI) for F-actin and nuclei visualization, respectively. Fluorescence microscopy was used to visualize cellular morphology. Confocal microscopy imaging, using 40X (*A*) and 80X objective (*B*), of human neutrophils treated with dimethyl sulfoxide (DMSO), N-formylmethionine-leucyl phenylalanine (fMLF), or GPR97 TA peptidomimetic (P7 and P18) for 10 minutes. *B*, Columns show separate images of biological triplicates. Red arrows indicate regions of polarized F-actin. Scale bars are 20 μm.

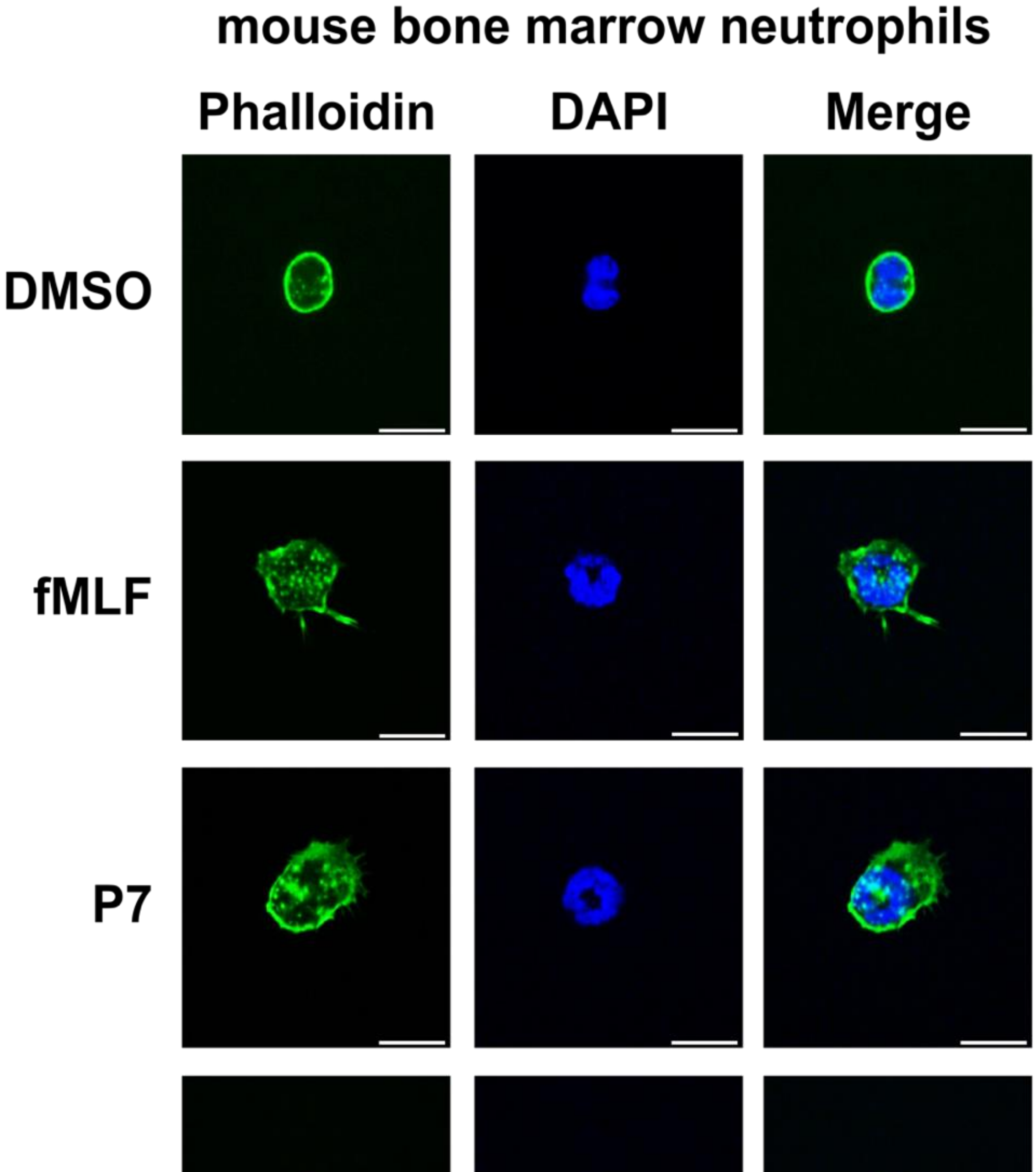


**Supplemental Figure 5: GPR97AP-dependent mouse bone marrow neutrophil polarization.** *A*, mBMNs were stained with phalloidin and 4', 6'-diamidino-2-phenylindole (DAPI) for F-actin and nuclei visualization, respectively. Fluorescence microscopy was used to visualize cellular morphology. Confocal microscopy imaging, using 100X objective, of mBMNs were treated with dimethyl sulfoxide (DMSO), N-formylmethionine leucyl-phenylalanine (fMLF), or GPR97 TA peptidomimetic (P7 and P18) for 10 minutes. Scale bars are 10 µm.

### AUTHOR CONTRIBUTIONS

T.F.B., F.K., Y.F., R.G., N.C., A.V.S., and G.G.T. designed and performed experiments. M.H. provided human blood and advice on neutrophil isolations. C.P. advised on neutrophil chemotaxis. A.V.S. advised on neutrophil signaling and conducted pilot studies. G.G.T. oversaw the direction of the research. T.F.B. and G.G.T. wrote the manuscript.


### CONFLICTS OF INTEREST

The authors declare that they have no conflicts of interest with the contents of this article.


### DATA AVAILABILITY STATEMENT

All the data are contained within the manuscript.
